# Consumers’ active choice behavior promotes coevolutionary units

**DOI:** 10.1101/2021.06.04.446921

**Authors:** Allan Maurício Sanches Baptista de Alvarenga, Marcelo Eduardo Borges, Leonardo Ré Jorge, Isabela Galarda Varassin, Sabrina Borges Lino Araújo

## Abstract

Individual behavior and local context are processes that can influence the structure and evolution of ecological interactions. In trophic interactions, consumers can increase their fitness by actively choosing resources that enhance their chances of exploring them successfully. Upon searching for potential resources, they are able to decide which one to choose according to their fitness benefit and maneuverability. Mathematical modeling is often employed in theoretical studies to understand the coevolutionary dynamics between these species. However, they often disregard the individual consumer behavior since the complexity of these systems usually requires simplifying assumptions about interaction details. Using an individual-based model, we model a community of several species that interact antagonistically. The trait of each individual is modeled explicitly and is subjected to the interaction pressure. In addition, consumers can actively choose the resources that guarantee greater fitness. We show that active consumer choice can generate coevolutionary units over time. It means that the traits of both consumers and resources converge into multiple groups with similar traits, exerting reciprocal selective pressure between them. We also observed that network structure has a greater dependence on the parameter that delimits active consumer choice than on the intensity of selective pressure. Consequently, this parameter can closely match empirical networks. Thus, we consider that the inclusion of consumers’ active choice behavior in the models plays an important role in the ecological and evolutionary processes that structure these communities.

## Introduction

Ecological interactions build the architecture of biodiversity in biological communities [1]. In trophic interactions such as parasitism, parasitoidism, predation or herbivory, individuals of one trophic level (consumers) exploit individuals of the trophic level below, as food resources. Consequently, these interactions result in increased consumer fitness at the expense of resource fitness. A foraging consumer will generally encounter different kinds of resources and they can decide which one to choose according to some ‘currency’ of biological fitness (e.g., rate of net energy intake, handling time, predator avoidance) [2,3]. This decision-making process known as ‘active predator choice’, leads the consumers to use some resources more often than others, given an encounter with each type of resource [4], e.g., birds that typically eat mollusks of particular sizes or species [5]; nest parasites that use the host’s nests whose eggs are similar to their own [6–8]; insects that differ in their oviposition patterns based on plant defense traits [9–12]; prey choice by hematophagous insects [13] or parasitoid insects that choose their prey through chemical signals [14].

Little is known about the evolutionary effects of adaptive diet choice on the dynamics and composition of ecological communities [15]. Theoretical studies on active consumer choice have been restricted to population dynamics, not considering its effect on community evolution [16,17]. However, ecological and evolutionary processes can be combined via natural selection [18] and occur on contemporary scales [19]. These eco-evolutionary dynamics, such as the relationship between the ecology of populations, communities and the evolution of functional traits, generates information that would not be expected in isolation [20]. The outcomes of eco-evolutionary dynamics between antagonistic species are generally related to the strength of selection imposed by the interaction [21,22]. The modeling of the active choice is simplified by assuming a random choice behavior combined with another function that determines the probability of interaction to occur successfully, depending on the adjustment of traits between consumer and resource [22,23]. This assumption implies that the consumer does not evaluate the resource’s trait, which increases the chances that it interacts with a resource that results in small fitness despite the presence of better resources available in its neighborhood. Such simplification may be understood as equivalent to active choice behavior since the imposed probability function will favor those interactions with a higher probability of success. However, this simplification does not limit the trait range that a consumer will try to interact. A first theoretical step addressing the effect of an active choice on species evolution was made for pairs of antagonistic interacting species [24], where it was observed that active consumer choice has evolutionary consequences. One of them, for example, is an unexpectable pattern where the resource trait is locked in only one of two evolutionary stable trait solutions [24]. Nevertheless, a theoretical framework investigating the effects of active consumer choice on coevolutionary dynamics in communities remains unknown.

A huge effort has been made to understand the mechanisms that determine the structure of interaction networks in communities [22,23,25–28]. Divergent selection regimes, phylogenetic conservation [29,30], habitat heterogeneity [31] and morphological attributes [32] may lead to nonrandom patterns of interactions and in the tendency of different subsets of species in the network to interact more frequently with each other than with the remaining species in the network [29,33–35]. Modularity play fundamental roles in ecological community resilience [36] and persistence since disturbances are not easily spread to other modules [37]. Besides that, modules have been suggested to be candidates for coevolutionary units [25,34]. That means that the modules are formed by coevolution and stay stable over time. However, it is not clear to date how such convergence could emerge in antagonistic networks, where the resource species selection pressure should tend towards divergence, not convergence.

Here, we integrate individual-based modeling with ecological networks tools to move forward our understanding of the role of the individuals’ active choice behavior in antagonistic network evolution. Our results demonstrate that the active consumer choice is a crucial element in giving rise to and promoting the stability of modules, generating coevolutionary units.

## Methods

### The model

We simulate an ecological system of two trophic levels that interact antagonistically, composed of several species and individuals that are explicitly modelled. Consumer attack traits and resource defense traits are subject to selection and mutation. The interactions occur through trait matching, that is, the probability of a successful interaction increases with the adjustment between the traits of the interacting individuals. A closer adjustment between both species traits is advantageous for the consumer and detrimental for the resource. Consumers actively choose resources within an interaction neighborhood, which represents the possibility of the consumer to evaluate the resources near them and choose which one will be attacked. In addition to the interaction pressure, we consider a stabilizing external pressure that models all types of pressure outside the interaction. This pressure acts as a selective force on consumer traits and resources towards a favored trait. Both the pressure of the interaction and the stabilizing pressure result in the fitness of the individuals, i.e., the contribution of these individuals to the next generation.

The model considers *M*_*X*_ resource species with *N*_*X*_ individuals per species and *M*_*Y*_ consumer species with *N*_*Y*_ individuals per species. It assumes the existence of a set of characters that constitute the defense or attack traits of individuals. Such characters may be morphological, physiological, chemical or behavioral and are represented by a real number, 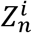, where *Z* represents the defense (*X*) or attack (*Y*) trait, *i* the individual and *n* the species. For example, 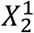 indicates the defense trait of individual 1 belonging to species 2 and 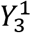 indicates the attack trait of individual 1 belonging to species 3.

### Dynamics

The dynamics of the model consists of three main steps in the following order: (i) the encounter between individuals; (ii) the fitness due to the interaction pressure and stabilizing pressure; and (iii) the reproduction (Fig.1).

**Fig. 1.**
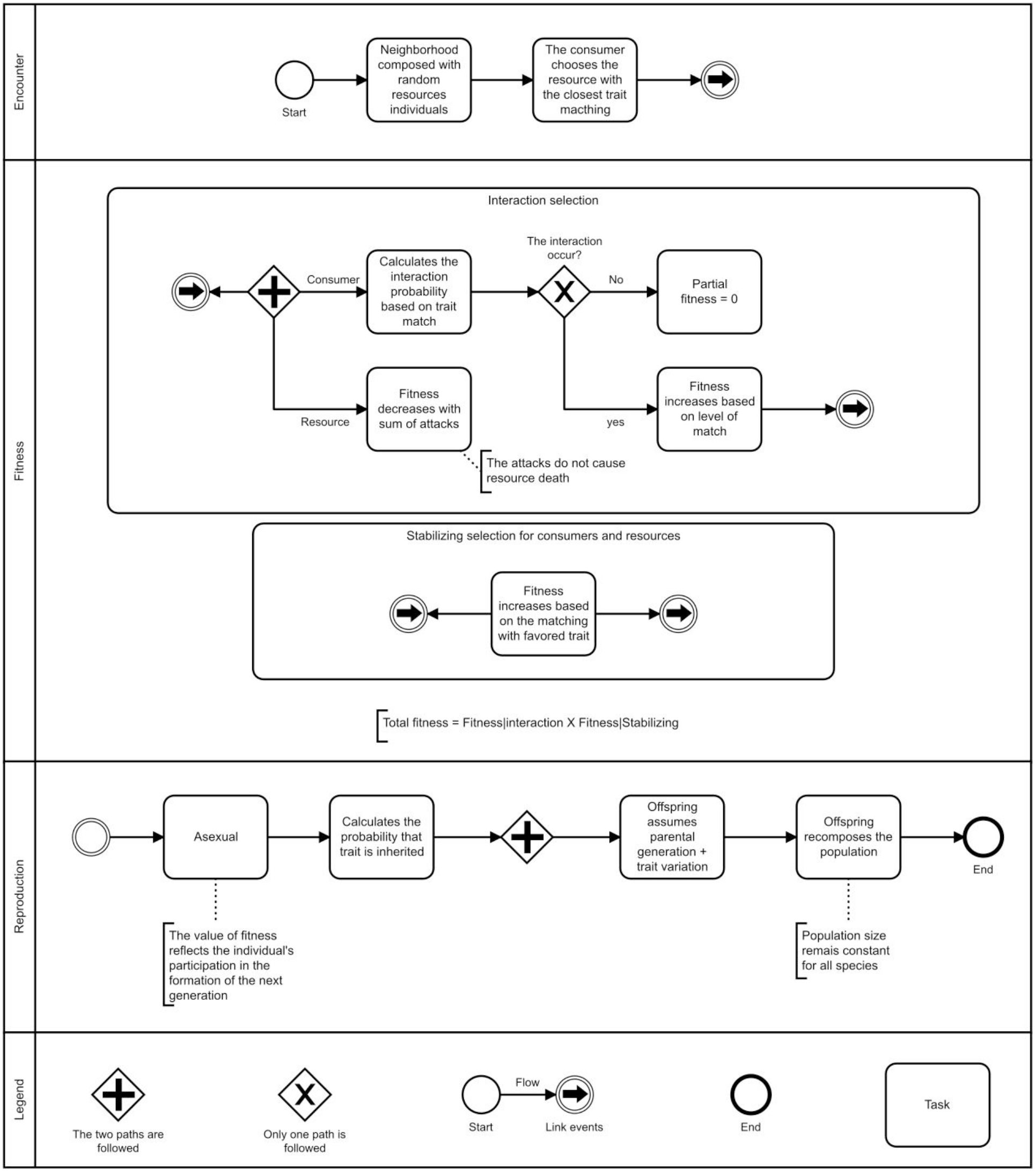
Steps of the model. The dynamics start with the encounter between consumers and resources within an interaction neighborhood. Consumer actively chooses and tries to interact with the resource that maximizes its fitness. Both consumers and resources have their total fitness calculated, composed of the partial fitness due to the interaction and stabilizing pressures. The result of the total fitness is reflected in the individual’s participation to the next generation.

#### (i) Encounters

#### (ii) Fitness 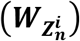

The total fitness of a resource individual 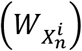 or a consumer individual 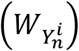 is given by the product of the performance of its trait due to the interaction and the selective pressure given by the external stabilizing selection:

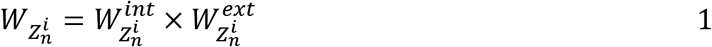

where, *Z* ∈ *X,Y*. The details of both selective pressures are detailed below:

##### Interaction pressure

We model the interaction mechanism based on trait matching, where the probability that the interaction occurs successfully increases as the difference of the consumer trait on resource decreases, according to:

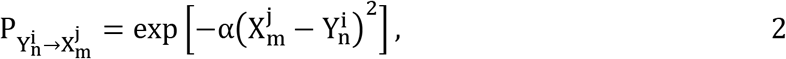

where *α* is a parameter that controls the intensity of the selective pressure on the interaction (Fig. S1a).

When an interaction occurs successfully, the consumer’s fitness due to the interaction also depends on the matching. Hence, if the interaction occurs successfully, a consumer’s fitness due to the interaction is given by:

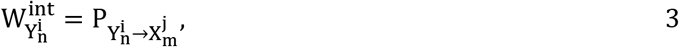

and if the interaction does not occur,

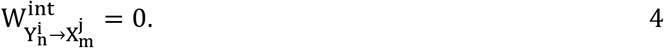

For the resource, both the intensity and number of attacks contribute to a decrease in its fitness. The attacks do not directly imply the death of the resource, but rather a decrease in its fitness:

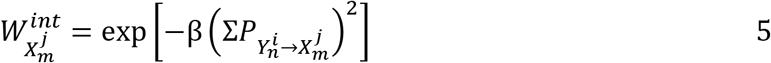

where *β* is a parameter that controls the intensity of the interaction pressure on the resource. A higher value of *β* penalizes resources whose phenotypic compatibility with the consumer is high, as it increases the impact of the attack of a consumer with high phenotypic compatibility with the resource (Fig. S1b). The term 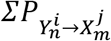 Eq.(5),represents the sum of all successful attacks weighted by the consumers’ interaction fitness. It means that a consumer that possesses greater trait matching will cause more impact on the consumer’s fitness than a consumer with smaller trait matching.

##### Stabilizing pressure

We include a stabilizing selective pressure, which considers all types of pressure outside the interaction and acts as a selective force on traits towards a favored trait, both in resources and consumers:

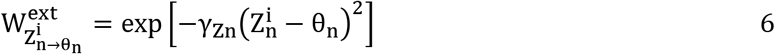

where *θ_n_* it is the trait favored by the external stabilizing selective pressure for a given species n and *γ* it is a parameter that controls the intensity of the pressure to the deviations of *θ*_*n*_. For simplicity we assume γ constant over species and trophic levels.

#### Reproduction

We assume that all individuals with non-zero fitness can have offspring which will then recompose the population to its original size. Thus, the number of individuals is constant over time, regardless of the number surviving a given generation. Our analyzes consider only those cases in which there was no extinction as, given these dynamics, extinction events occur only in extreme situations. Therefore, the participation of the individual *i* to the next generation is proportional to its fitness relative to other individuals of the same species:

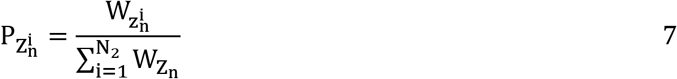

where 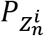 it is the probability that an individual of the new generation will inherit the trait 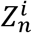 of the individual *i* of the *n* species. 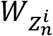 refers to the fitness of the parental individual Eq.(7), and 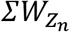 the sum of the adaptive values of all individuals of the parental species.

For simplicity, the reproduction is asexual and the offspring assumes the same trait value as the parental individual with a mutation coefficient δ, whose value is a random number that follows a normal distribution probability:

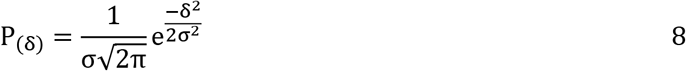

where *σ* is the standard deviation, which we assume constant between trophic levels.

##### Simulation parameters

In all the simulations the number of species, the number of consumer and resource individuals per species, and the intensity of external stabilizing pressure were maintained constant (*M*_*X*_ = 50, *M*_*Y*_ = 50; *N*_*X*_ = 100, *N*_*Y*_ = 100, *γ* = 1, respectively). The traits favored by the stabilizing selection of the resource and resource species were obtained from a normal distribution *θ* ∼ *N*(0,1) (mean equal to 0 and a standard deviation equal to 1). Therefore, the simulated community presented heterogeneity of trait values favored by the external stabilizing selection.

We ran simulations without active consumer choice under different intensities of interaction pressures (see values of α and *β* in Table 1) to verify their effect on the Coevolutionary trait dynamics. The model without this behaviour is obtained by assuming the interaction neighborhood is equal to a single resource individual, which corresponds to *Φ* = 0.02%. In simulations with active consumer choice, the intensity of interaction pressure was fixed (*α* = 0.8 and *β* = 0.2). These two values correspond to intermediate values approached in the case without active choice. Also, different sizes of the interaction neighbourhoods *Φ* were evaluated. All the values of parameters and variables used in the simulations are described in Table 1. Each simulation consisted of 10,000 generations. To verify the model’s sensitivity to random events, five replicates of each simulation were performed (146,491 networks in model with active choice and 5,145 in the model without active choice). The simulations were carried out in FORTRAN language both in the LCPAD - Central High-Performance Processing Laboratory, Federal University of Paraná and through the Amazon web service and will be available online.

**Table 1.** Parameters used in the simulations, their values and a short description

| Parameter/variable | Value | Description |
| --- | --- | --- |
| $M_X, M_Y$ | 50, 50 | Number of species of resources and consumers. |
| $N_X, N_Y$ | 100, 100 | Number of resource and consumer individuals by species |
| $\delta$ | random number that follows a normal distribution probability which standard deviation is $\sigma$ | Mutation coefficient |
| $\sigma$ | 0.02 | The standard deviation used to calculate phenotypic variation due to reproduction |
| $\gamma$ | 1 | Stabilizing pressure intensity |
| $\theta$ | $\theta \sim \mathcal{N}(0,1)$ for consumers and resources | Trait favoured by stabilizing pressure |
| $\alpha$ | 0, 0.05, 0.1, 0.2, 0.4, 0.8†, 1.6, 3.2, 6.4 | Intensity of interaction pressure on the consumer |
| $\beta$ | 0, 0.05, 0.1, 0.2†, 0.4, 0.8, 1.6, 3.2, 6.4, 12.8, 25.6, 51.2, 102.4 | Intensity of the interaction pressure on the resource |
| $\Phi$ | from 0.02% to 20% an increase by 1% and from 20.2% to 100% an increase by 10% | Size of the interaction neighbourhood (0.02 % implies that the attack is without active choice) |

† represents values of $\alpha$ e $\beta$ that were kept constant in simulations that the effects of variation in the interaction neighborhood size was investigated.

### Data analysis

#### Interaction persistence networks

To evaluate the persistence of interactions over time we built an interaction persistence network, from the matrix of size *N*_*x*_ × *N*_*y*_, where each row and column represent a resource and a consumer species, respectively. The value of each cell indicates the number of generations in which at least one interaction between the given pair of species was recorded. To avoid transient effects, we only used the data for the last 4,000 generations, sampled at every 200 generations, resulting in 21 networks per simulation. We have also analyzed the interaction network for each time, where each cell of the matrix represents the number of interactions between a pair of species (see Supplementary Material).

The interaction persistence networks were characterized using established network metrics: connectance (*C*), modularity (*M*) [38] and specialization index (*H2’*)[39]. The measure of all the mentioned metrics was implemented through the bipartite package and performed in an R [40] environment. Modularity was measured using the DIRTLPAwb+ algorithm using the *computeModules* function [41]. Specialization *H2’* was measured using the *H2fun* function [41]. Both metrics use quantitative matrices. The connectance (*C*) was calculated in binary matrices and refers to the ratio between the number of non-zero cells by the matrix size [41]. The connectance indicates the percentage of all interaction occurred during the analyzed time. Higher values of modularity in the interaction persistence networks indicate that species interactions occur more often (in time) in a subgroup of species than between them. Similarly, the higher the specialization index, it means that a pair of species persists their interaction over time more intensely than expected by the abundance of species.

## Results

In most cases, active consumer choice led to coevolutionary trait dynamics with stable groups of tightly interacting species that exert reciprocal selection on traits. Within each module, the resource traits converge into a narrow range of values, surrounded by consumer traits (Fig. 2a and Fig.S2). Smaller neighborhoods induced more extreme trait dynamics, with average trait values reaching double the amplitudes of larger neighborhoods (see Fig. 2a: *Φ* = 4%). In these smaller neighborhoods, there was a high frequency of interactions (darker colors in Fig 2b and Fig. S3) between consumers and resources within each module over generations. That is, all species interact with each other inside the modules in most generations. In larger neighborhoods, due to a higher opportunity of encounters with preferred resources, the frequency of interactions over generations between all species inside the module decreased (lighter colors). However, the presence of interactions highlights that, among the species in the same module, a consumer species changes its choice of interaction over time. This alternation is maintained over the generations, but it is locked inside the module without breaking the unit of coevolution. Stable coevolutionary units were not observed in scenarios without active consumer choice (Fig 2c, Fig S4) and the interactions occur between almost all species regardless of the interaction pressure intensity (Fig. 2d). Additionally, we observed that even higher interaction pressure intensities do not promote coevolutionary units, but instead drive species to extinction (Fig. S4).

**Fig. 2.**
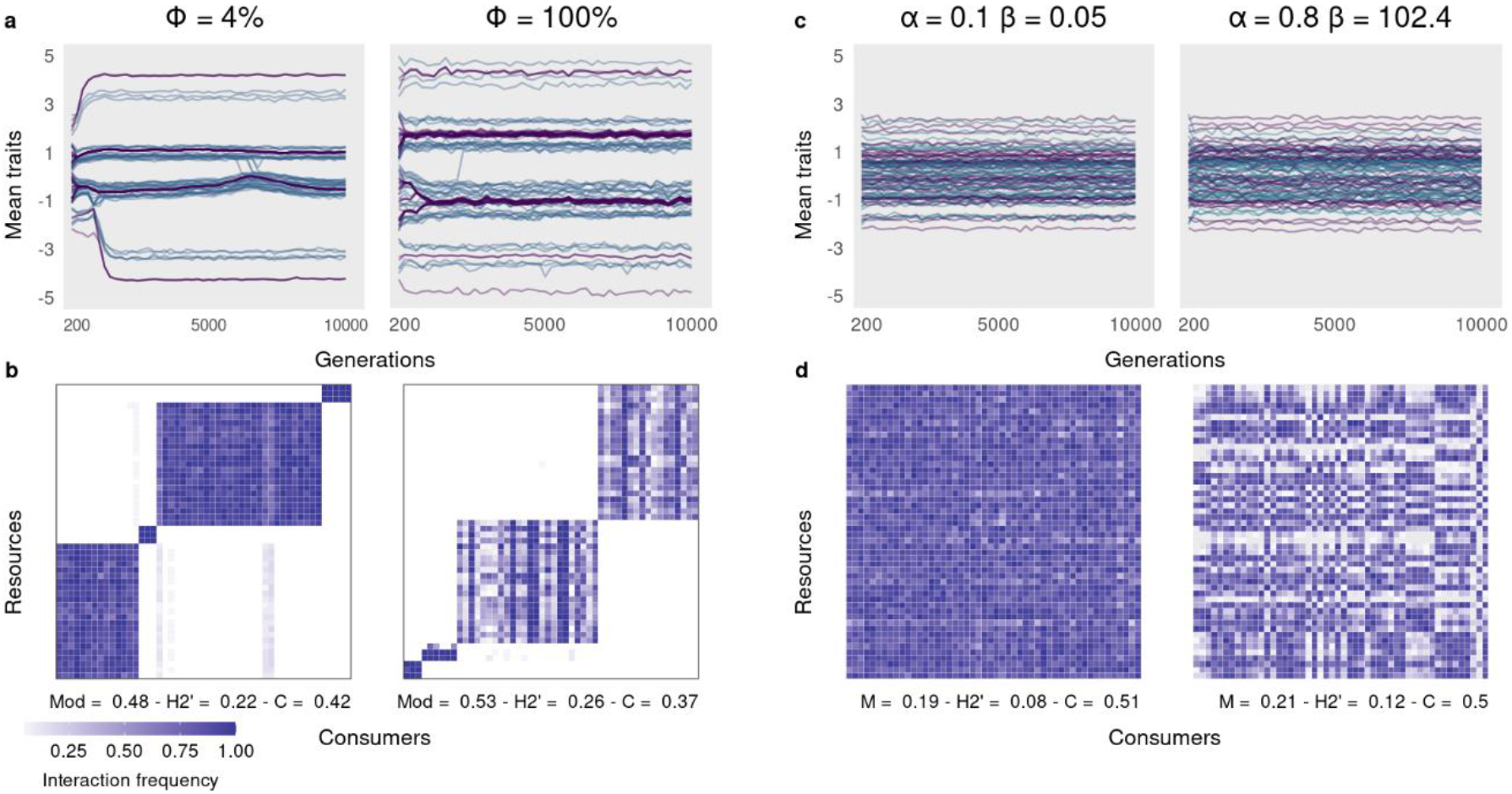
Coevolution under and without active consumer choice. **(a-b)** Coevolutionary trait dynamics under active choice in different sizes of interaction neighborhoods. (c-d) Coevolutionary trait dynamics without active choice behavior, but under different interaction pressures. Figures (a) and (c) show the average trait of each consumer species (blue) and each resource species (purple). Figures (b) and (d) represent the matrices of interaction persistence: the frequency of generations in which at least one interaction between a pair of species was recorded. The absence of interaction is represented by the color white. Network metrics: M = Modularity; H2’ = Specialization; C = Connectance; Note that active choice behavior limits species interactions to subgroups, evidencing the stability of the evolutionary units.

As the size of the interaction neighborhood increases, the network tends to be more modular, specialized, less connected and consumer success decreases (Fig S6). However, for *Φ* between approximately 0.2% and 1%, this trend is inverted for all metrics. This inversion occurs when the first coevolutionary units emerge, but with only two or three modules, which increases the interactions between species, explaining the metric inversions (Figs S2 and S6). For *Φ* around 1% and higher, the metrics follow the initial trend again (Fig S6). However, between approximately 1% and 10%, the coevolutionary units oscillate between two and four modules, varying both over time and over replicates. For Φ around 10% and higher, the coevolutionary units stabilize (Figs. S2 and S3 and Fig 2). To avoid this initial variation, we restrict the next results to *Φ* > 10.

The network metrics showed considerable difference according to consumer choice behaviour. Without active choice but varying the interaction pressure (α and *β*) connectance ranges from 0.47 to 0.51; modularity from 0.17 to 0.22; and specialization, from 0.08 to 0.14. With active choice, and *Φ* > 10, connectance ranges from 0.29 to 0.48; modularity from 0.41 to 0.65 and specialization, from 0.19 to 0.32. Then, networks with active consumer choice were more modular, more specialized, less connected, and with lower consumer success in relation to networks without active choice behaviour (Fig. 3).

**Fig. 3.**
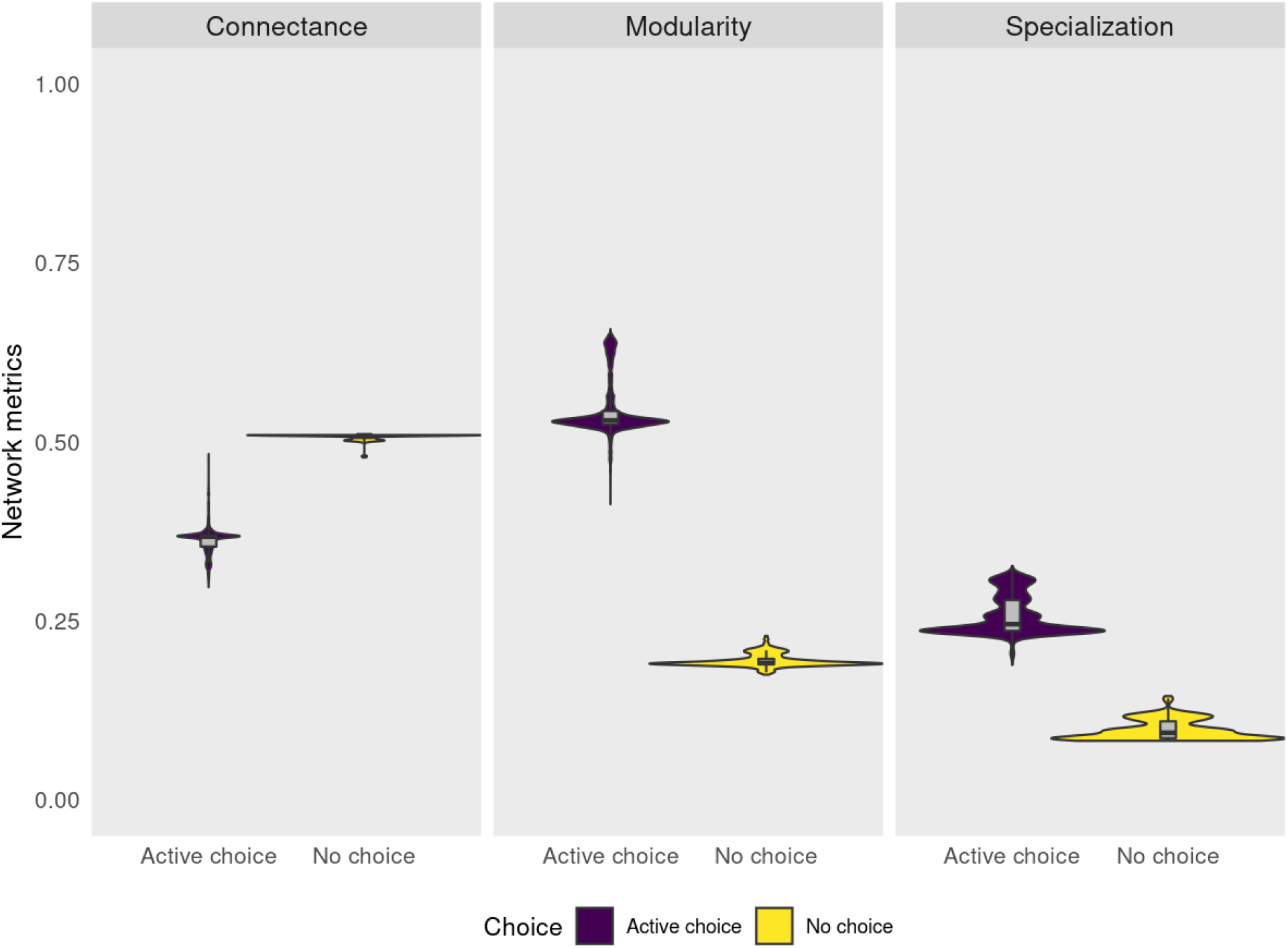
Network metrics with and without active choice behavior. Active choice promotes networks with lower connectance and higher modularity and specialization. The purple color indicates the model with active choice, the yellow color indicates the model without active choice. The violin plot shows the distribution of the data and the boxplots presents the summary statistics median and interquartile ranges.

## Discussion

In this study, we investigated the role of individual active choice behavior on coevolutionary trait dynamics and antagonistic network structure of species with antagonistic interactions. Our results reveal that active choice can drive significant changes in trait distributions, on the selective regimes and on patterns of interactions that shape the structure and dynamics of antagonistic networks. We demonstrate that the active choice behavior generates modules that are persistent in evolutionary time, which can be interpreted as co-evolutionary units. These results highlight the importance of individual behavior and the effects of adaptive diet choice on eco-evolutionary dynamics.

The model with active choice behavior allows each consumer individual to choose to interact with the resource in its neighborhood that maximizes its fitness. The simulations showed that, under this condition, subgroups of resource species converge their traits around a single value, while subgroups of consumer species converge their traits around one of two values - below or above resource traits values - locking the resource trait evolution (Fig.2a). These subgroups of resource and consumer species then form a temporal stable module with almost no interaction between modules. The mechanism behind this stability is probably the same observed for the model with two species [24]. That study analytically showed that active choice behavior locks the resource trait because any variant resource that maximizes consumer fitness will not go unnoticed by the consumer. Here, a small variation in a resource trait makes it a better resource choice by any of the surrounding consumers, reducing the resource fitness. On the other hand, without active choice, small variations in the resource trait are more likely to go unnoticed by the consumer, so that the temporal stability of the module as well the convergence of species traits are broken (Fig. 2b). Thus, higher pressure intensity on the interaction (α and *β*) is not a sufficient ingredient to increase modularity and trait convergence.

Modules have been suggested to be candidates for coevolutionary units [22,25,34], implying that modules are formed by coevolution and stay stable over time. Inside the modules, the species interact with each other, exerting strong reciprocal selection on traits, shaped by a similar regime of selective pressures [25,31]. Andreazzi et al. (2017) proposed a model for antagonistic interactions, and observed that coevolutionary units can emerge from antagonistic interaction, but only when the fitness consequence is higher for the consumers than victims. Here we show that active choice implies a higher increase in modularity and stability than in the models without active choice (Fig 3). Since the enlargement of the interaction neighborhood increases the fitness consequence for the resources, our model supports that evolutionary units can emerge even under high pressure on resources.

Coevolutionary units have been suggested as a product of cospeciation and arms race. Under These hypotheses, the consumers are predicted to have evolutionary patterns of diversification that are congruent with the patterns of their resources, where closely related resource species would have similar defenses and closely related consumers would feed on closely related resources [42,43]. However, these hypothesis has little support in empirical studies [42,44–47], except for tight specialized interactions [48]. The incongruence between host and parasite phylogenies, for example, has previously been explained in terms of host switching, extinction, duplication events and failure of the parasite to speciate in response to host speciation [49]. In fact, our model does not predict interaction only between pairs of species, which would be the first step of cospeciation. We show that the coevolutionary units in antagonistic interactions also produce convergent traits, independently of cospeciation (or any speciation, as our model has static species), and even when the consumer can choose among all resources (*Φ* = 100), species interact with almost all other species within the module. Further studies must be done to investigate if diversification patterns could emerge from our model.

The mechanism behind the coevolutionary units may be the emergence of convergent traits among individuals of the same trophic level, for example the presence of mainly white flowers inside the module in mutualistic interactions networks ([34]; [25]; [31]). This arises due to the reciprocal fitness benefit among the two trophic levels, which does not occur in antagonistic interactions, and thus trait convergence is not expected. However, it has been observed that where distantly related plant species share a common assemblage of herbivores, they are likely to defend themselves with similar strategies [50]. Besides, consumers experience a selection pressure to evolve specific traits adapted to consuming the existing resource species [51] that is, they “track” resource defenses and not resource species per se [43]. For example, closely related herbivores prefer Inga (tree) hosts with similar defenses rather than closely related Inga [52]. Regardless of these examples, there is not yet a mechanistic explanation on why distant related resources would converge their traits since they could develop different strategies to defend themselves. Our results suggest that resource trait convergence promotes attack dilution: when resources converge their traits, the pool of options for a consumer increases and the chance of a specific individual being attacked decreases. In other words, with different resource species with similar phenotypes, the effects of the attacks of the consumers are diluted among the resources inside the module.

In this study, we were able to evaluate the effects of active choice behavior in eco-evolutionary dynamics using simplifications (see methods). We suggest that future studies include more ingredients in modeling to capture more information about this mechanism. For example: (i) asexual inheritance can be a limitation for generalization of the model, although there are several types of antagonistic interaction in which the interacting species present asexual reproduction, as in interactions between bacteria and viruses [53–57] or bacteria and protists [58,59], bacteria and nematodes [60,61], daphnias and parasites [62,63]; (ii) the spatial homogeneity disregards the differences between landscapes, as well as gene flow limitations [31,64]; (iii) although the model has an evolutionary time scale, it does not allow speciation, which could reveal how the individual behavior can promote species diversification; (iv) finally, the equivalence between generations of consumers and resources disregards differences in consumer and resource life spans, when it is common to have several generations of consumers in relation to a single generation of the resource, as in parasite-host relationships [34].

To conclude, we show that consumer active choice of resources that maximize their fitness is a crucial element for the emergence of coevolutionary units, that is, modules formed through the coevolutionary process. Moreover, as far as we know, this work is the first to demonstrate the mechanism of dilution by which traits converge in antagonistic networks.

## Data accessibility

R-project to reproduce the full paper and FORTRAN input: Allan Maurício Sanches Baptista de Alvarenga, Marcelo Eduardo Borges, Leonardo Ré Jorge, Isabela Galarda Varassin, & Sabrina Borges Lino Araújo. (2021, March 17). Alvarenga et al. 2021 data set and project (Version 1). Zenodo. http://doi.org/10.5281/zenodo.4606671

## Authors’ contributions

SBLA conceived the idea of the project, SBLA, AMSBA and MEB wrote the model and performed the analyzes. AMSBA wrote the R code for the data analyses and AMSBA, MEB, LRJ, IGV and SBLA interpreted the results and contributed to the writing of the manuscript.

