## Supporting information for "Consumers’ active choice behavior promotes coevolutionary units"

November 20, 2020

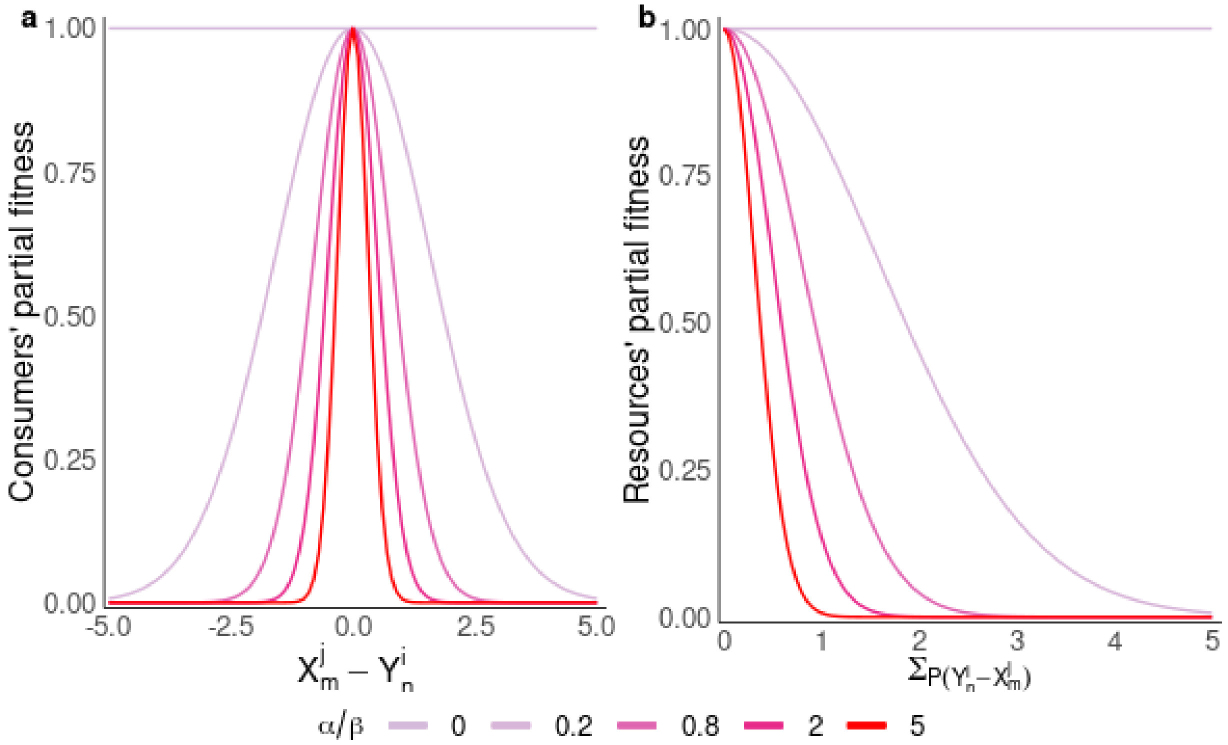


Fig. S 1: Fitness functions due to interaction. (a) Part of the consumer's partial fitness as a function of phenotypic differences between interacting individuals. (b) resource interaction pressure as a function of the intensity of the attacks (sum of all successful attacks weighted by the consumers' interaction fitness (Eq. 2.5)). The lines represent distinct values of the intensity of the selective pressure on the consumer ($\alpha$) and on the resource ($\beta$) used in the simulations.

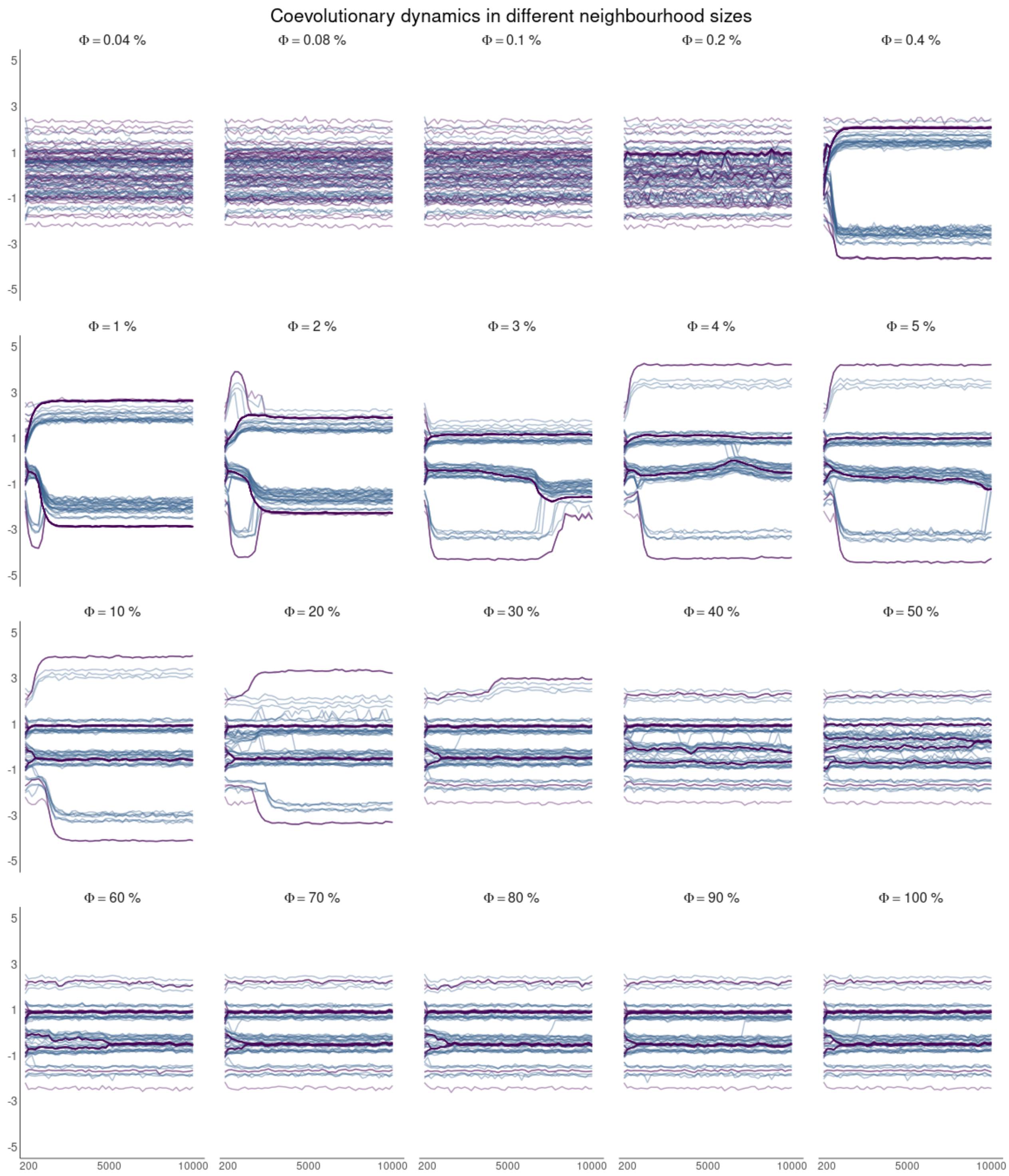


Fig. S 2: Coevolutionary trait dynamics in different neighbourhood sizes of interactions in the model with active choice. The average traits of the consumer species are represented by the blue lines and the resource species are represented by the purple lines. Note that in intermediate values of $\Phi$ the traits evolve to more extreme values. The active choice in conjunction with the size of the interaction neighbourhood changed the coevolutionary trait dynamics. The dynamics were similar to the scenario without active choice in neighborhoods with sizes between $\Phi = 0.04\%$ and $\Phi = 0.2\%$. From that value of $\Phi$, the traits of the resources converged. These converging traits evolved towards the ends of the phenotypic space, moving away from the optimal values imposed ($\theta_i$) by the stabilizing selection. Consumers also showed a convergence of traits (see $\Phi = 0.4\%$, for example). In even larger neighbourhoods, resource traits were trapped between consumer groups with similar traits, forming coevolutionary units (see $\Phi = 100\%$, for example).

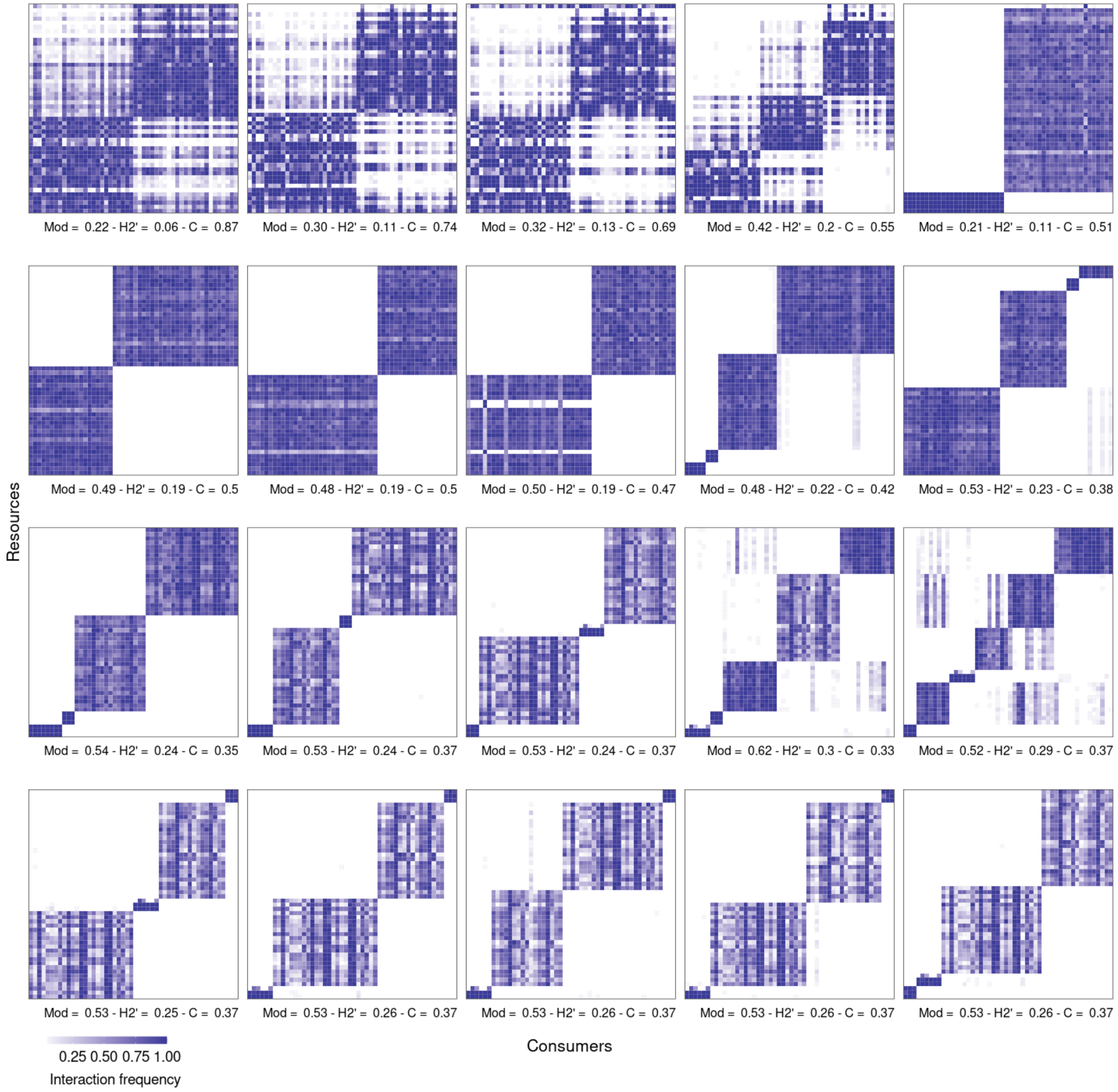


Fig. S 3: Matrices of interaction persistence under active choice in different sizes of interaction neighbourhoods. The frequency of generations in which at least one interaction between a pair of species was recorded. The absence of interaction is represented by the colour white. Network metrics: Mod = Modularity; H2' = Specialization; C = Connectance; Note that active choice behaviour limits species interactions to subgroups, evidencing the stability of the evolutionary units. Note that In smaller neighbourhoods, there was a high frequency of interactions (darker colours) between consumers and resources within each module over generations, that is, all species interact with each other inside the modules in most generations. In larger neighbourhoods, due to a higher opportunity of encounters with best resources, the frequency of interactions over generations between all species inside the module decreased (lighter colours).

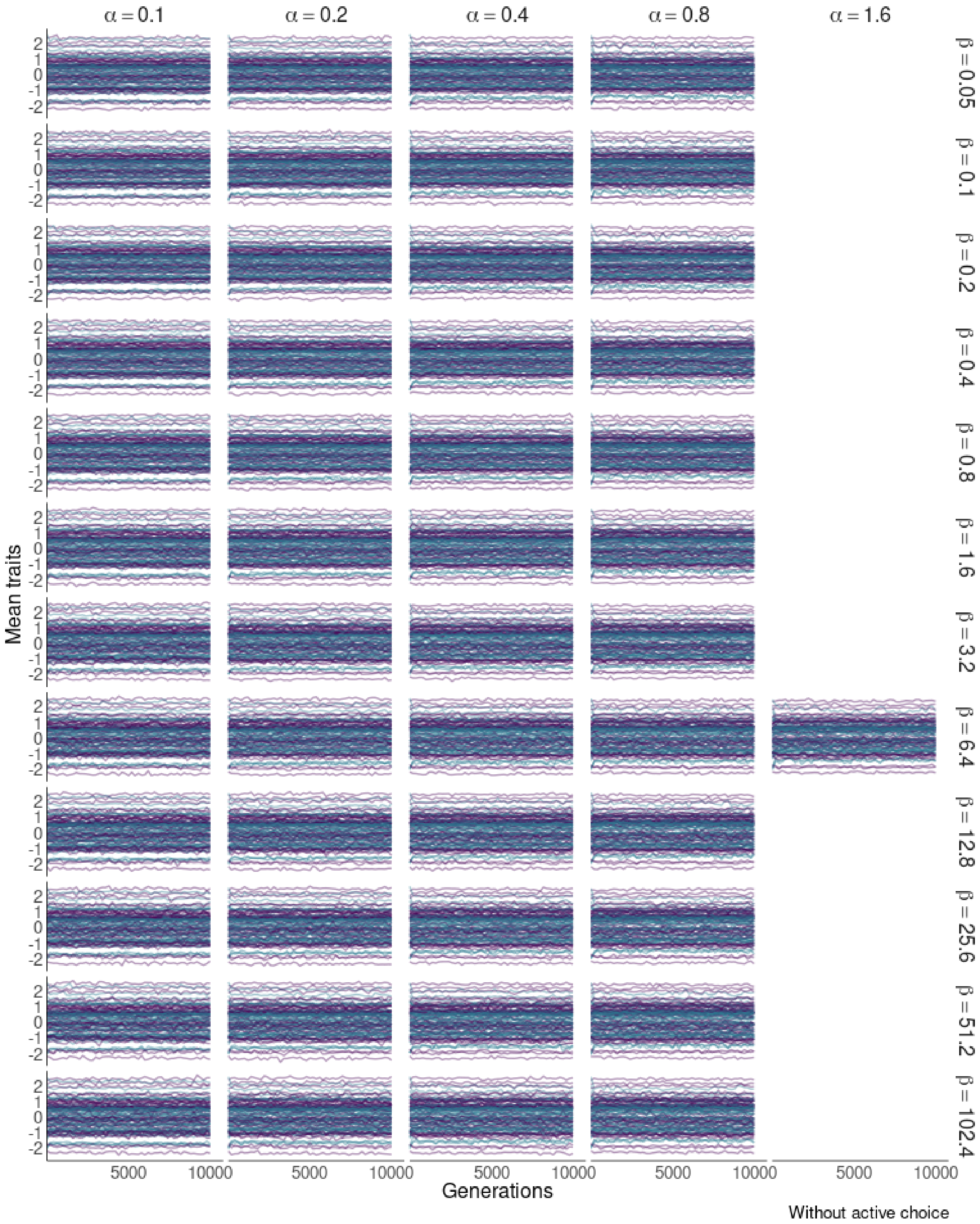


Fig. S 4: Coevolutionary trait dynamics in different intensities of interaction pressure in the model without active choice. The average traits of the consumer species are represented by the blue lines and the resource species are represented by the purple lines. Note that there is no convergence of phenotypes and that with higher intensities of selective pressure on the consumer lead to cases of extinctions (blank areas).

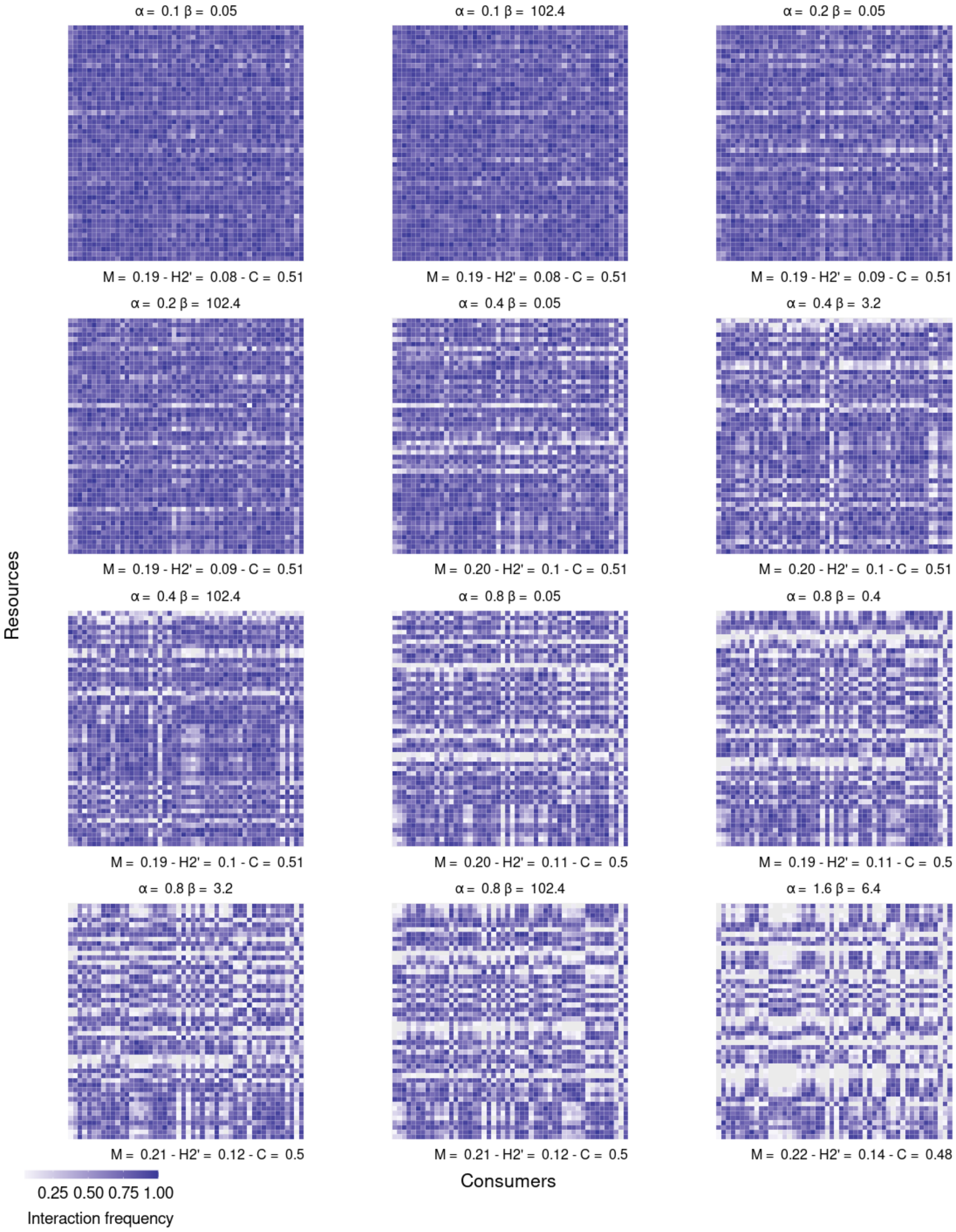


Fig. S 5: Matrices of interaction persistence under different selective pressures. The frequency of generations in which at least one interaction between a pair of species was recorded. The absence of interaction is represented by the colour white. Network metrics: Mod = Modularity (M); H2' = Specialization; C = Connectance; Note that there is no formation of isolated modules in cases without active choice regardless of the intensity of the selective pressure.

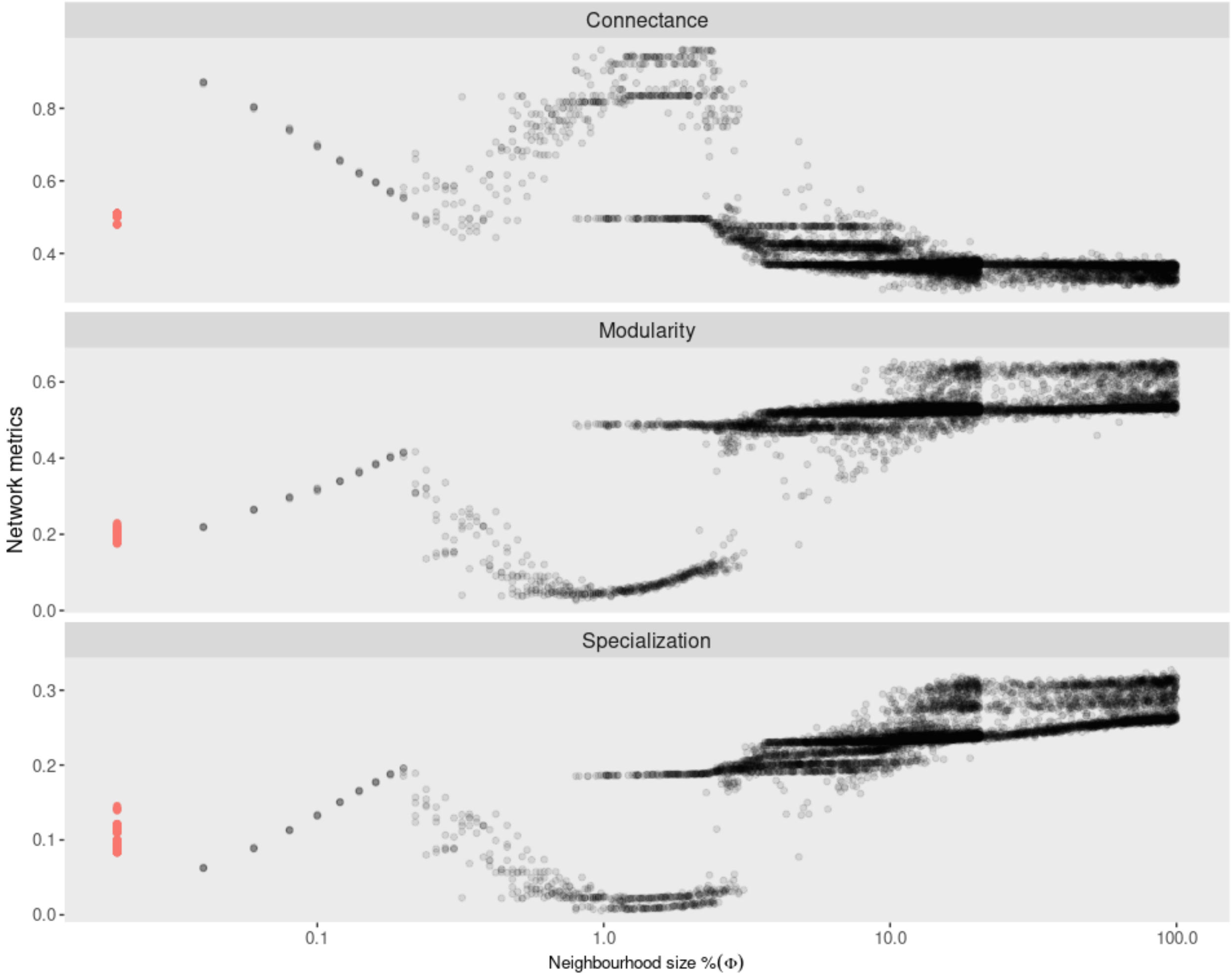


Fig. S 6: Network metrics in different sizes of interaction neighbourhoods.The red dots highlight the cases where the neighborhood is minimal (just one individual), which is equivalent to the scenario without active choice. As the size of the interaction neighborhood increases, the network tends to be more modular, specialized, less connected and the consumer success decreases. However, for $\Phi$ between approximately 0.22% and 1%, this trend is inverted for all metrics, then it returns to the first trend for $\Phi > 1\%$ .
